# Integrative structural interactomics reveals protein organization and structure in a giant virus

**DOI:** 10.1101/2025.06.16.659922

**Authors:** Lars Mühlberg, Julia Ruta, Vasilii Mikirtumov, Raymond Burton-Smith, Kazuyoshi Murata, Mikhail Kudryashev, Kenta Okamoto, Boris Bogdanow, Fan Liu

## Abstract

Giant viruses are large DNA viruses that infect unicellular and multicellular eukaryotes and form exceptionally large extracellular particles. (Meta)genomics and (meta)transcriptomics have provided insight into their diverse coding repertoire, but many of the proteins remain to be characterized as they lack homology with known proteins. Here, we integrated cross-linking mass spectrometry, quantitative proteomics, computational tools and cryo-EM data to characterize the protein architecture of intact melbournevirus particles. Based on this, we allocated 88 viral proteins to different virion sub-compartments and proposed topologies of 25 inner membrane proteins. We assigned eight components of the capsid in cryo-EM data, including proteins that tether the capsid shell to the membrane, reflecting key points in virion maturation. The data provide a valuable resource and demonstrate the power of an integrative approach to gain system-level structural insights into a poorly characterized biological system.

## Introduction

Advances in (meta)genomics and (meta)transcriptomics have revealed the remarkable genetic diversity of viruses, the largest on earth ^1–3^. More recently, breakthroughs in protein structure prediction, such as Alphafold2/3, have enabled large-scale modeling of viral protein structures ^4,5^. However, approximately 40% of the viral proteins lack known structural homologs ^6^. Moreover, the performance and accuracy of AlphaFold’s predictions depend heavily on the availability of sequence data ^7^, which is limited for many viruses, especially emerging ones ^8^. These challenges hinder our ability to infer protein structures and functions solely through *in silico* analyses, underscoring the need for complementary experimental approaches to study poorly characterized viruses and their interactions with the host.

Giant viruses are a prime example of complex yet poorly understood biological systems. As members of the nucleocytoplasmic large DNA viruses (NCLDV), they infect unicellular and multicellular eukaryotes and produce very large extracellular viral particles ^9^. Their particle size (ranging from 200 nm to up to 2 µm) and genome content (up to 2.8 megabase pairs) challenge conventional definitions of viruses ^10–12^. Intriguingly, giant viruses encode proteins involved in processes once thought exclusive to cellular life, such as metabolism, protein synthesis, or genome condensation ^13^. For example, the giant melbournevirus was found to encode three histone-like proteins (MEL_368, MEL_369 and MEL_149) that compact its viral DNA - a process previously associated primarily with eukaryotes ^14,15^.

Melbournevirus, first described in 2014, belongs to the *Marseilleviridae* family and has a highly complex, 360 kbp-long genome containing 403 predicted open reading frames (ORFs) ^16^. The virus replicates and assembles in amoeba *Acanthamoeba castellanii*, and newly formed virions exit the host cell via exocytosis or cell lysis ^16–19^.

Despite its significance, the structure of melbournevirus remains poorly characterized. The particle has a diameter of 230 nm and consists of several components: (1) an outer capsid shell ^20^, formed by homotrimeric major capsid protein (MCP, MEL_305), along with penton and cap proteins ^21^, (2) an internal membrane layer enclosing the viral DNA genome, and (3) a minor capsid protein (mCP) layer beneath the MCP layer, likely stabilizing the capsid structure and linking it to the inner membrane, a feature also observed in other NCLDV capsids ^22–25^.

The architecture of these complexes has been determined exclusively by cryo electron microscopy (cryo-EM) ^20–24^. However, the large size of capsid imposes resolution limitations that can be achieved in cryo-EM studies ^22^. Due to these constraints and the complexity of the system, only three Melbournevirus proteins have been structurally characterized: the major capsid protein MEL_305, and two nucleosome-like proteins MEL_368 and MEL_369 ^14,26^. Additionally, many melbournevirus ORFs are predicted to encode proteins lacking available homology to any known proteins in databases ^10^, severely constraining our understanding of the identity, structure, function, interactions, and spatial organization of the viral proteins in the virion. These challenges could potentially be addressed by analyzing the virion’s native protein interaction networks. Cross-linking mass spectrometry (XL-MS) is particularly well-suited for this task, as it enables the mapping of protein interaction networks within intact cells ^27^, organelles ^28^ and virions ^29^. Furthermore, integrating XL-MS with computational protein structural prediction offers a powerful approach to validate and refine structural models ^30–32^.

In this study, we combined proteome-wide XL-MS with AlphaFold3-based structure prediction to characterize the structural interactome of intact melbournevirus particles. Our XL-MS data enabled spatial mapping of 88 viral proteins to the two virion sublayers. Notably, within the membrane separating these two sublayers, we identified 25 viral transmembrane (TM) proteins and determined their topologies. Curiously, most viral TM proteins position the majority of their non-TM domains within the capsid-membrane space. By combining XL-MS, AlphaFold3 and cryo-EM data, we demonstrate that two of these TM proteins structurally bridge the inner virion membrane and the capsid, functioning as minor capsid proteins (mCPs). Furthermore, we generated structural predictions for multiple uncharacterized viral proteins, proposed molecular functions based on their structure, localization and interaction partners and assigned previously unresolved cryo-EM densities. Our findings provide detailed structural insight into melbournevirus protein organization and establish integrative structural interactomics as a powerful approach for multi-level analysis of giant viruses. More broadly, this methodology represents an attractive strategy for studying poorly understood biological systems.

## Results

### The melbournevirus interactome clusters packaged proteins to sub-virion compartments

We employed XL-MS to uncover protein-protein interactions (PPIs) within the large and poorly characterized melbournevirus particles. Virions were collected from the supernatant of infected *A. castellanii* cell culture and cross-linked using the MS-cleavable cross-linker disuccinimidyl sulfoxide (DSSO). To enrich virions and deplete contaminating debris, we purified the particles by sucrose density-gradient ultracentrifugation, as described previously ^16,20^. Proteins were then extracted, digested by trypsin, and cross-linked peptides were identified by LC-MS (Figure 1A).

**Figure 1.**
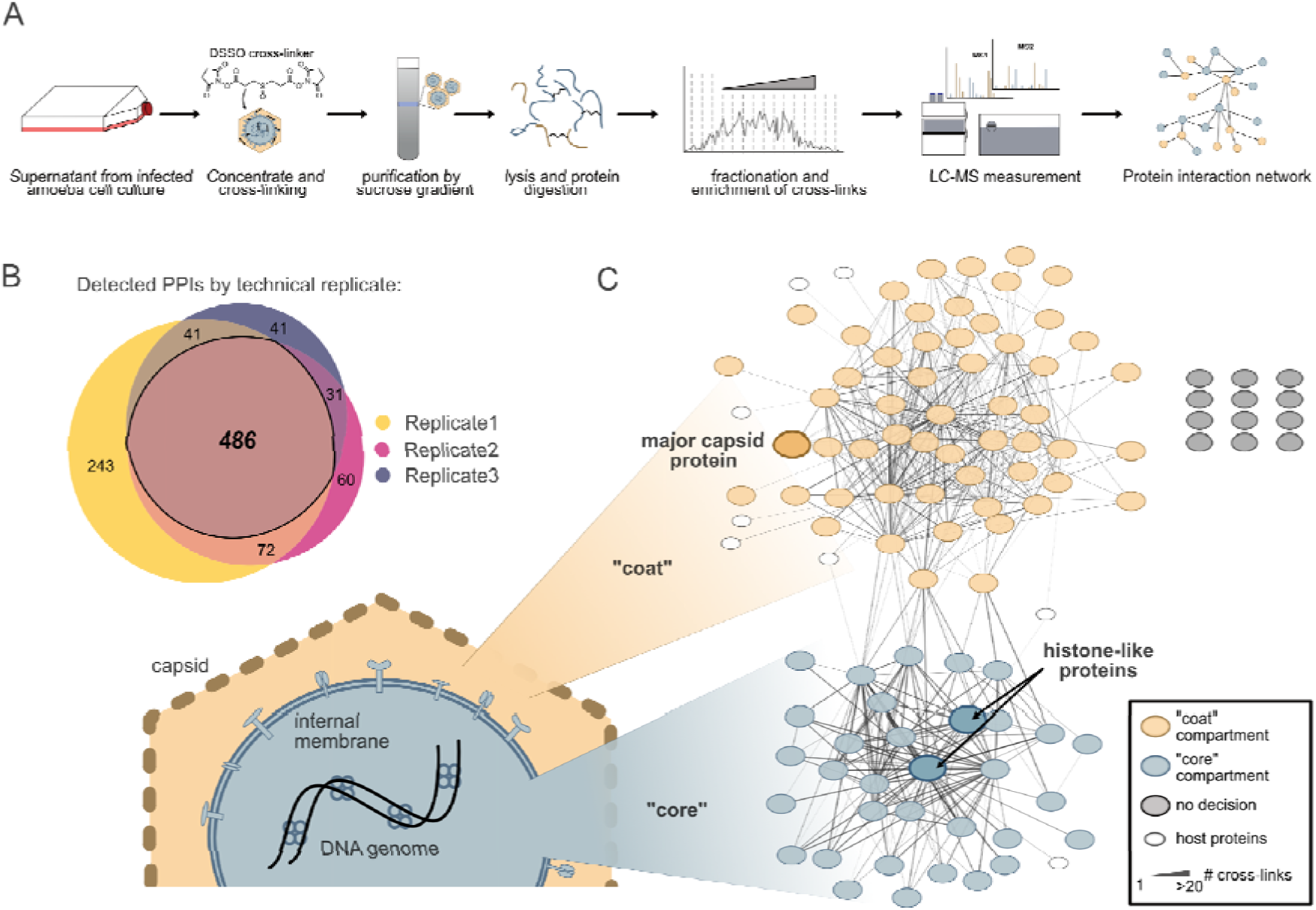
Protein interactome profiling of melbournevirus by XL-MS. **(A)** Workflow for XL-MS-based PPI profiling of intact melbournevirus particles. **(B)** Venn diagram of PPIs detected in three biological replicates. PPIs were aggregated from cross-linked peptide pairs filtered at 1% FDR. **(C)** XL-MS-based interaction network of melbournevirus. Proteins are categorized by localization: “coat” (external to the membrane, yellow), and “core” (internal to the membrane, blue). Proteins with only intra-protein cross-links are colored gray. The final dataset comprised 8,908 cross-linked peptide pairs, with PPIs retained only if detected in all replicates. Line thickness corresponds to the number of cross-links supporting each PPI. The network with annotated gene names is shown in supplementary Figure 1.

To assess the reproducibility of our approach, we performed XL-MS across three biological replicates. We identified between 4,900 and 8,100 unique residue-to-residue connections at a residue-level false discovery rate (FDR) of 1% (supplementary Table 1). After aggregating cross-links to PPIs, we analyzed the overlap between replicates. Approximately 64% of PPIs were identified in at least two replicates and 50% were present in all three, indicating high experimental reproducibility (Figure 1B). To minimize false positives, we retained only PPIs observed in at least two replicates. The resulting filtered interaction network comprised 486 PPIs, corresponding to 8,908 cross-links from 108 proteins (Figure 1C, supplementary Figure 1, supplementary Table 1).

While most cross-links involved viral proteins, a small fraction (2 - 4.5% per replicate) connected viral and host proteins. We included only those Amoeba host proteins with direct viral interaction partners, likely representing virion-recruited host proteins.

These included amoebal Cystatin, an HMG (High mobility group) box domain-containing protein, Peptidyl-prolyl cis-trans isomerase, Vinculin and several unannotated proteins corresponding to previously identified transcripts in Amoeba ^33^. The predominance of viral proteins and the scarcity of highly abundant host proteins inside the virion align with prior proteomic analyses ^34^.

Next, we investigated the spatial distribution of packaged proteins. Previous cryo-EM studies revealed that melbournevirus possesses a 30 □-thick inner membrane layer separating the outer capsid from the inner dsDNA genome and associated proteins ^20^. This membrane imposes a constraint on XL-MS, as the DSSO cross-linker, which has a spacer arm of 10.1 □, cannot connect proteins on opposite sides. To evaluate this constraint in our PPI network, we performed unsupervised community clustering ^35^, which partitioned the network into two major protein communities (supplementary Table 2). Proteins within the same community exhibited dense interactions, whereas connections between communities were sparse. These communities reflect spatial segregation of a genome-associated internal “core” and a membrane-external, capsid-enclosed “coat”.

We then predicted sub-virion localizations for melbournevirus proteins based on the only four proteins with known localizations: the histone-like viral proteins (MEL_368 [H4-H3-analog], MEL_369 [H2B-H2A-analog] and MEL_149 [“mini” H2B-H2A-analog] ^14^ localized to the core, while the major capsid protein (MCP, MEL_305) resided in the coat ^20,21^. Using the spatial information of their interaction partners, we classified the viral proteome into 33 core and 55 coat proteins. This represents the first system-wide spatial mapping within a giant virus.

### Transmembrane predictions and cross-linking data assign topologies to viral TM proteins

Next, we sought to identify proteins bridging the two communities, focusing on membrane-anchored proteins in our cross-linking dataset. Since no experimental data were available for transmembrane (TM) domain annotation of melbournevirus, we screened the viral proteome using *in silico* prediction tools (DeepTMHMM, DeepTMpred and TMbed; Supplementary Table 3). These algorithms predict both α-helical and β-barrel TM regions with high accuracy ^36–39^. To enhance reliability, we retained only TM domains predicted by at least two algorithms. While TMbed and DeepTMHMM showed strong agreement, DeepTMpred assigned more TM domains (Figure 2A), likely because it does not distinguish between TM domains and signal peptides ^37,39^. This analysis identified 43 putative TM proteins, of which 25 were present in our interaction network (Figure 2A).

**Figure 2.**
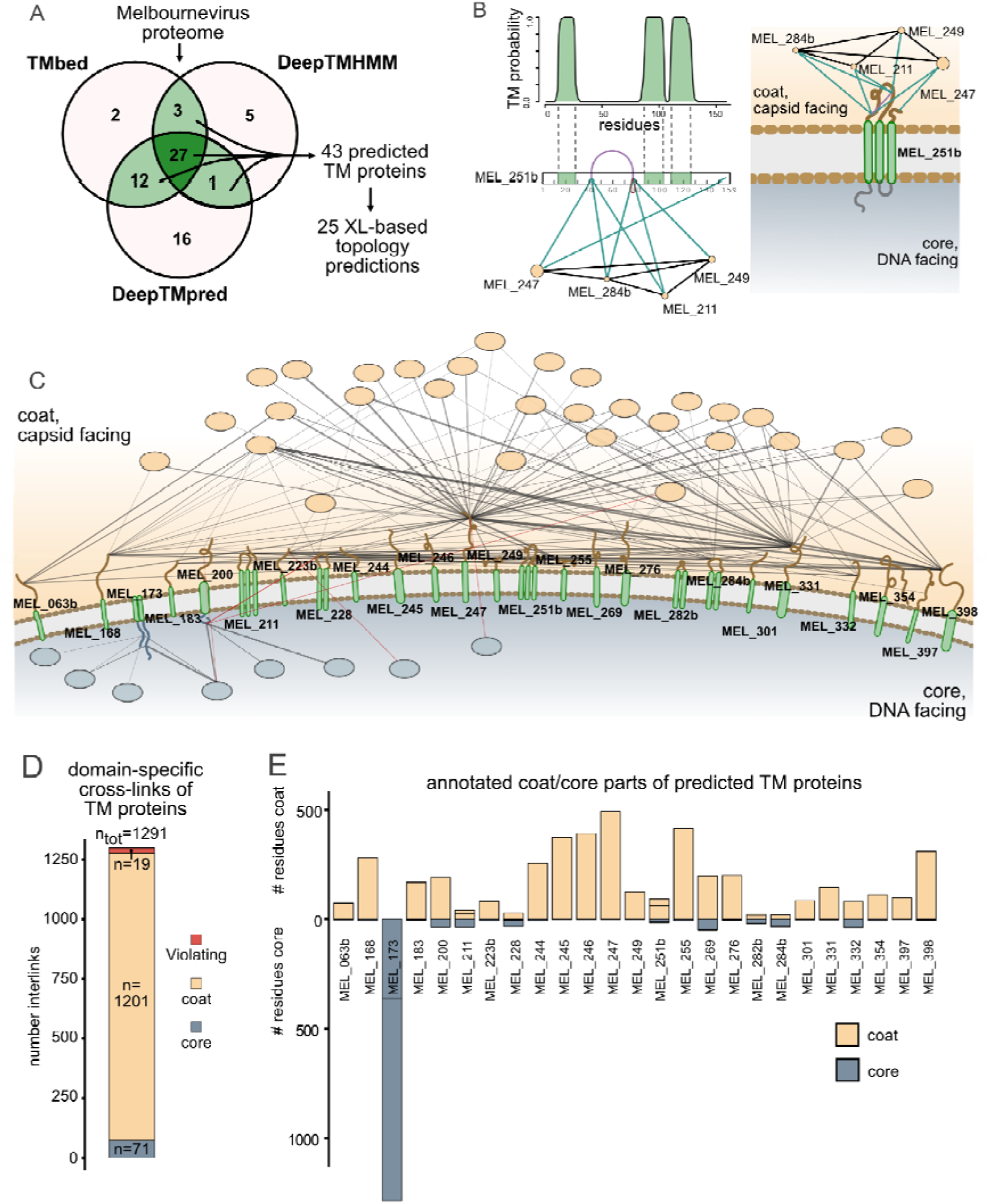
Topology analysis of virion-incorporated transmembrane proteins in melbournevirus. **(A)** Venn diagram of the TM proteins predicted by three different algorithms within the melbournevirus proteome. Only TM proteins with detected intra-protein cross-links were included in further analysis. **(B)** Representative example of topology annotation, combining predicted TM domains (green) and cross-linking data. Regions that cannot be predicted (e.g., lacking cross-links) are shown in gray. Interacting proteins are color-coded by predicted localization as in Figure 1. **(C)** Overview of predicted TM protein topologies and their direct interaction partners. Cross-links within the same domain are grouped for clarity and gene names of interactors from coat and core compartments are removed. Cross-links supporting the proposed topology are shown in black; those violating the topology are highlighted in red. A fully annotated network is shown in supplementary Figure 2. **(D)** Number of cross-links in the three different categories shown in panel C. **(E)** Number of amino acids in the “coat” and the “core” of the 25 predicted TM proteins. In case of multiple domains per protein, the domains in the same compartment are added onto another as indicated by black lines separating those parts.

A key advantage of XL-MS is its domain-level resolution, enabling topology prediction of TM proteins ^28^. Leveraging this, we assigned topologies based on the localizations of interacting partners in the PPI network (Figure 2B). Using 1,291 cross-links, we predicted topological orientations for 29 domains of 25 predicted TM proteins (Figure 2C, supplementary Figure 2, supplementary Table 3). These predictions explained 98.5% of the cross-linking data (Figure 2D). The remaining ∼ 1.5% of cross-links conflicted with our predictions, possibly due to mis-incorporated proteins, false identifications or proteins with dual topologies ^40^.

Notably, most cross-links on TM proteins mapped to the outer membrane leaflet. Consistently, non-TM domains were predominantly located in the coat (Figure 2E). This suggests melbournevirus TM proteins evolved with large coat-facing domains and small core-facing regions. One exception is MEL_173, which has two non-TM domains, both predicted in the core. Taken together, our XL-based analysis resolved a high-resolution sub-virion interactome, revealing dozens of viral TM proteins with a preferential outer-leaflet exposure.

### Cross-links validate structure prediction of melbournevirus proteins

Having established the global melbournevirus interactome with sub-virion resolution, we next pursued atomistic structural insights. We predicted structures for all viral proteins in our cross-linking network (Figure 3A) using AlphaFold3, focusing on proteins containing at least one intra-link (cross-links connecting lysines within the same protein sequence). Model quality was assessed using predicted template modeling (pTM) scores ^5^.

**Figure 3.**
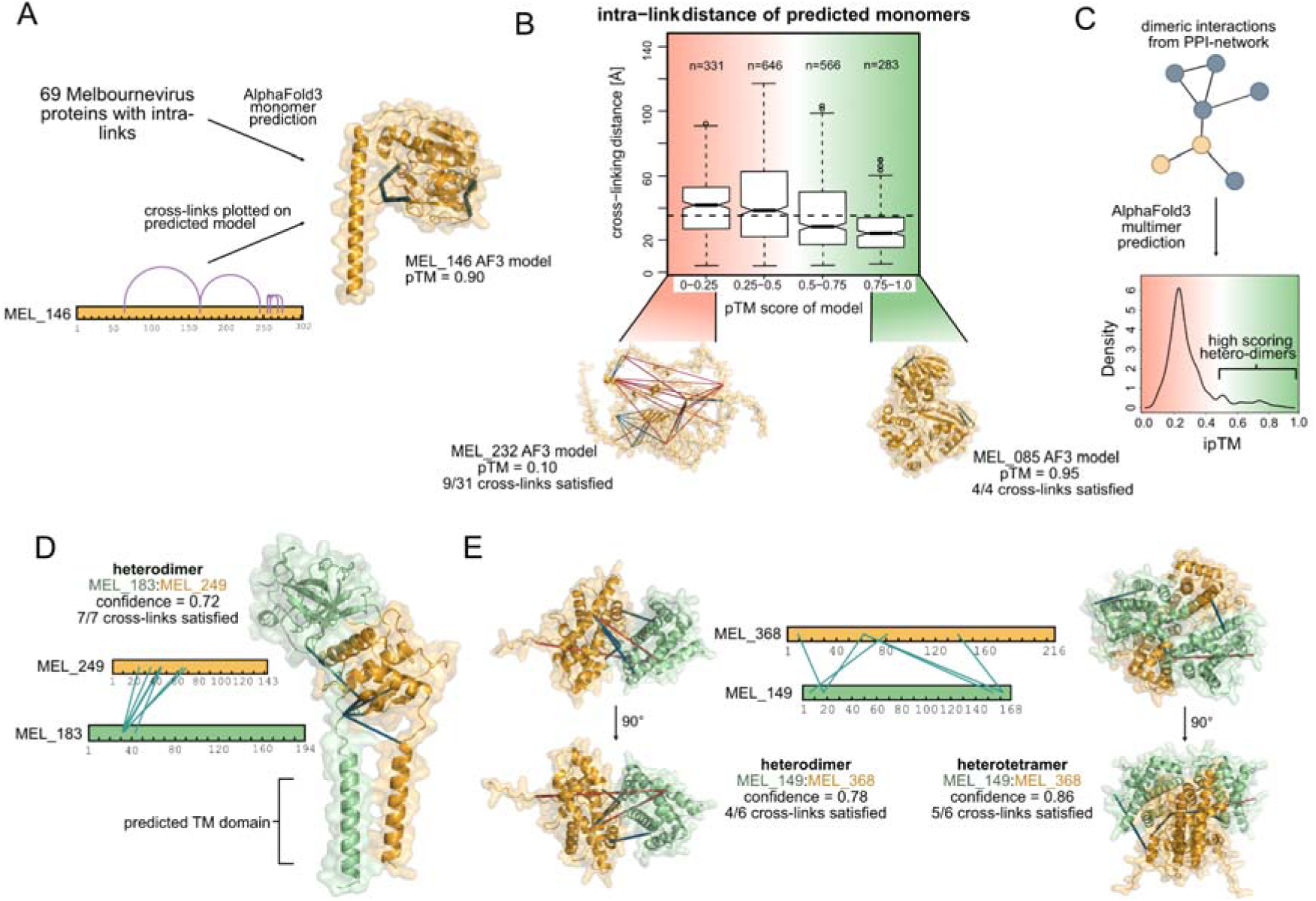
AlphaFold3-based structure prediction and cross-linking validation. **(A)** AlphaFold3-predicted structures of 58 Melbourneviral proteins with intra-protein cross-links, evaluated using a 35 Å maximum distance constraint. Representative example (MEL_146) shows cross-link mapping (blue lines) on the predicted structure. **(B)** Agreement between cross-link-derived distances (Cα–Cα) and model confidence (pTM scores; very low, 0.2-0.4; low, 0.4-0.6; medium, 0.6-0.8; high 0.8-1.0). Higher-confidence models better satisfy the 35 Å constraint than low-confidence predictions. **(C)** Heterodimeric structural prediction of cross-linked protein pairs by AlphaFold Multimer. Density plot of interface pTM (ipTM) scores reveals limited high-confidence predictions (right-skewed distribution). **(D)** Heteromultimeric prediction of two TM proteins, MEL_183 and MEL_249. All inter-protein cross-links from structured regions comply with the 35 Å threshold. **(E)** Predicted structures of histone-like proteins MEL_368 (H4-H3 analog) and MEL_149 (H2B-H2A analog) as heterodimers and heterotetramers (stoichiometry inferred from prior knowledge of melbournevirus nucleosome data). Agreement of inter-protein cross-links and AlphaFold confidence score support the heterotetrameric model.

To validate these predictions, we mapped intra-protein cross-links onto the AlphaFold3 structures and measured Cα-Cα distances (supplementary Tab 3). Models with low pTM scores (e.g., MEL_232, pTM = 0.1) frequently violated the DSSO distance constraint of 35 □ ^41^, while high-scoring models (e.g., MEL_085, pTM = 0.95) showed excellent agreement with cross-linking data (Figure 3B). Models with a pTM score of 0.5 and higher generally agreed with the cross-linking data and can therefore be considered high-confidence predictions. Overall, AlphaFold3 generated 45 high-confidence structures where >50% of intra-links satisfied distance restraints (supplementary Table 4).

In addition to structures of individual proteins, we expanded our analysis to prediction of heterodimers with AlphaFold3 ^5^. After filtering by the AlphaFold confidence score, we identified 16 high-confidence dimeric models consistent with our cross-linking data (confidence score >0.5, at least 50% of inter-protein cross-links from structured regions satisfied, Figure 3C and supplementary Table 4). A notable example is the MEL_183:MEL_249 complex (confidence score = 0.72), where all seven inter-protein cross-links satisfied distance constraints (Figure 3D). This model is further supported by the observation that both proteins are membrane incorporated (see also Figure 2) and the globular domains building the interaction interface are both located in the “coat”.

Another insightful prediction is on the viral histone-like proteins. We obtained high-scoring heterodimeric models for MEL_368 (also known as the melbournevirus histone H4-H3-analog) with either MEL_369 (also known as the melbournevirus histone H2B-H2A-analog) or MEL_149 (also known as mini H2B-H2A-analog). While MEL_368:MEL_369 forms canonical nucleosome-like complexes within the virion, MEL_368 and MEL_149 association remained enigmatic ^14^ and were only recently observed to form a complex in vitro ^42^. Comparative analysis of heterodimer versus heterotetramer predictions revealed a slight improvement of the confidence scores (from 0.8 to 0.9) and increased cross-link concordance (from 3/5 to 4/5) (Figure 3E). These data support MEL_368:MEL_149 may form an alternative, minor nucleosome-like species which is present within native virions.

### High-confidence protein models pinpoint the putative functions of uncharacterized viral proteins

Leveraging our high-confidence structure models, we investigated potential functional roles of melbournevirus proteins through structural homology searches ^43^. Using Foldseek ^44^ we compared the 39 high-scoring, cross-link validated monomeric models against the PDB and AlphaFold databases. Remarkably, 25 models showed significant structural similarity (probability score > 0.95) to known protein folds (supplementary Table 5). Many melbournevirus proteins, especially in the coat, share structural similarities with enzymes. For example, MEL_085 displays a fold similar to amino acid oxidases, while MEL_146 and MEL_398 resemble glycosyltransferases (Figure 4A). In the core, many protein models showed structural similarity to proteins involved in DNA-related processes. For example, MEL_275 aligns with TATA-box binding proteins, MEL_380 is related to serine/threonine protein kinases, and MEL_219 has an RNase III-like structure.

**Figure 4.**
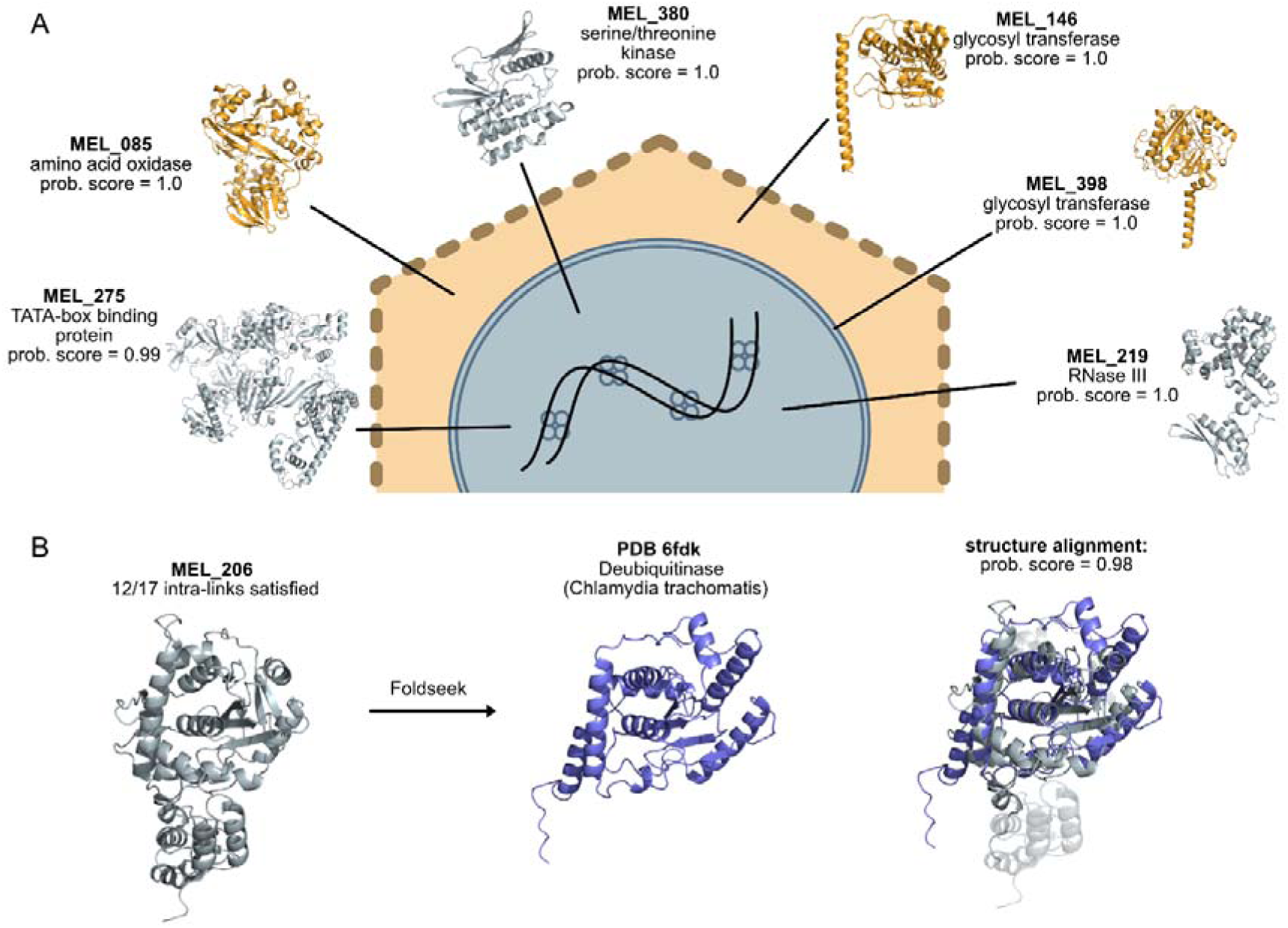
Structural comparison of high-confidence models with known PDB structures. **(A)** Structural similarity analysis using Foldseek, identifying matches between high-scoring melbournevirus protein models (probability score ≥0.95) and Protein Data Bank (PDB) entries. Selected hits are annotated with predicted molecular functions and virion localizations. **(B)** Representative structural alignment showing MEL_206 (gray) superimposed with a ubiquitin-specific protease from *Chlamydia trachomatis* (PDB: 6FDK ^70^, blue). Right panel displays probability scores comparing MEL_206’s predicted structure with PDB: 6FDK.

A particularly intriguing case is MEL_206 – a protein with no sequence homology to characterized proteins ^16^, but displaying striking structural similarity to several ubiquitin-specific proteases (Figure 4B). The model aligned closely with the structure of Chlamydia trachomatis DUB1 (deubiquitinase and deneddylase), suggesting potential ubiquitin protease activity. Supporting this hypothesis, we studied protein ubiquitination of the virion proteome by proteomics and found several ubiquitinated peptides originating from the histone-like proteins MEL_368 and MEL_369 (supplementary Table 6). Both proteins are cross-linking partners of MEL_206. These findings suggest MEL_206 may regulate viral chromatin accessibility through histone ubiquitination as a potential regulatory mechanism.

### Refining the cryo-EM capsid structure by integrative modeling

In addition to putative enzymes, we sought to further characterize the structural properties of melbournevirus proteins. Current cryo-EM reconstructions of melbournevirus particles have resolutions ranging from 4.4 □ ^21^ to 3.4 □ ^45^. While certain regions of the map permitted de novo model building, the complexity of the system, unknown stoichiometry, and limited resolution in many other regions made de novo modeling based solely on density unfeasible ^21,45^. To determine whether our integrative XL and structure modeling approach could facilitate structural analysis, we leveraged protein connectivity data from XL-MS to assign proteins in the 3.4 □ map ^45^.

We began by fitting MEL_305 (MCP), the only previously assigned capsid protein ^21^ into the map. Reassuringly, the predicted structure aligned well with the cryo-EM density, underscoring the accuracy of high-confident AlphaFold models (supplementary Figure 3).

Beneath the MCP layer (Figure 5A) is a supporting layer of minor capsid proteins forming a complex network. Among these, the pentasymmetron component (PC□) exhibits a rhodopsin-like stacked α-helical fold, though its identity remained unresolved in existing cryo-EM data ^21^. To identify this protein, we screened high-scoring models among predicted coat proteins and found that density matched to MEL_213b, when predicted as a homodimer. Furthermore, homodimeric prediction of MEL_213b satisfied the XL-distance restraints (supplementary Figure 3).

**Figure 5.**
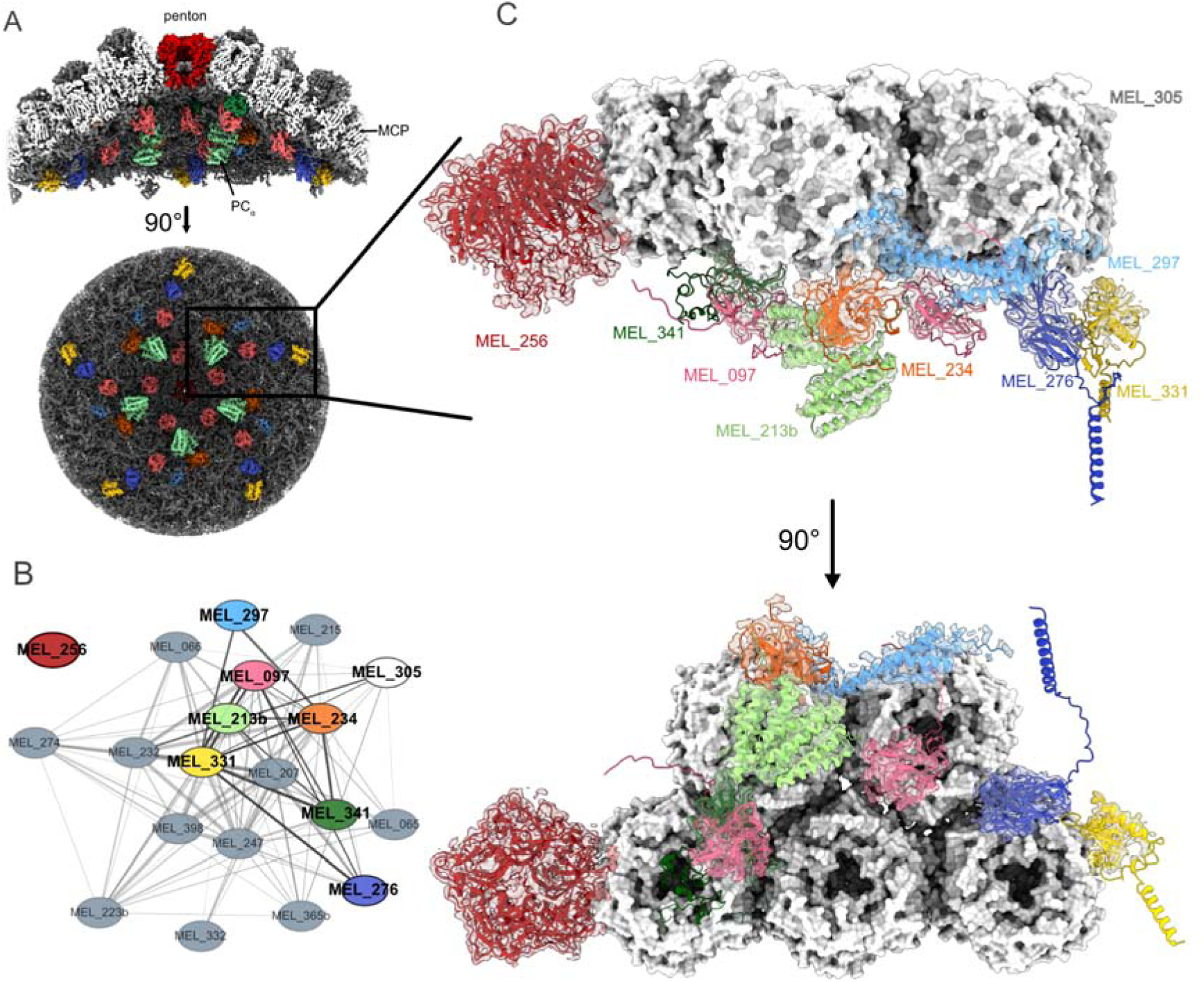
Structural annotation of the melbournevirus capsid using AlphaFold models and cryo-EM data. **(A)** The melbournevirus cryo-EM map focused on a five-fold symmetric capsid vertex (side and interior views, data from ^45^). Different components of the capsid are color-coded in ChimeraX. **(B)** Part of the PPI network of the protein community located in the coat compartment and connected to MEL_305 or MEL_213b. Proteins MEL_256 and MEL_297 have also been added for completeness. Identified capsid-associated proteins are color coded as in A. **(C)** Fitting of AlphaFold models into the cryo-EM density. Proteins are color coded the same as in panel A and B. Major capsid protein MEL_305 is shown in surface-mode for clarity.

To identify additional minor capsid components, we shortlisted candidate proteins from the coat community that cross-linked to either MCP or MEL_213b (Figure 5B). By fitting these models into the cryo-EM map, we assigned MEL_234 and MEL_276 as minor capsid proteins (Figure 5C). We also identified four other proteins - MEL_341, MEL_097, MEL_297 and MEL_331 - whose AlphaFold3 models contained well-structured domains consistent with the cryo-EM density. However, their lower accuracy domains could not be fitted, likely due to their high flexibility. In such cases, PPI data provided critical orthogonal evidence. Interestingly, MEL_276 and MEL_331 were predicted to be membrane associated (see also Figure 2C), suggesting a structural role in capsid-membrane tethering.

Each vertex of the MCP icosahedron is characterized by the presence of a homopentameric complex, the penton ^21^. Assigning the identity proved challenging, as none of the candidate models cross-linking to MCP or MEL_213b displayed a similar fold. We therefore screened all XL-MS detected proteins for potential homo-multimerization (n=1-5, supplementary Figure 4 and supplementary Table 7). As expected, MCP and MEL_213b scored the highest at their known stoichiometries. The only protein with a high-scoring pentameric model was MEL_256, which fit exceptionally well into the cryo-EM map (Figure 5C). The detected inter-domain cross-link further supported the MEL_256 pentamer (supplementary Figure 3). Refinement of the AlphaFold3 models by tracing revealed an extensive contact between the penton complex and the minor capsid protein MEL_341, which wraps around the vertex center beneath the penton (supplementary Figure 5). Remaining unassigned densities in the cryo-EM map lacked sufficient evidence for confident model matching (supplementary Figure 6).

To extend these findings, we assessed the stoichiometry and symmetry of assigned capsid components using shotgun proteomics and intensity-based absolute quantification (iBAQ) values from purified virion samples. By calibrating copy numbers against four reference proteins (with known abundances from literature or our cryo-EM assignments), we obtained highly reproducible estimates across four technical replicates (supplementary Figure 7). This analysis suggested ∼120 copies of MEL_097 and MEL_276, and ∼ 60 copies of MEL_297 and MEL_341 (supplementary Table 7). The other minor capsid proteins such as MEL_234 (∼2200 calculated copies) and MEL_331 (∼550 calculated copies) are incorporated in much higher stoichiometries and therefore are also likely part of the capsid outside the vertex (previously named “scaffold” component, “glue” or “zipper” ^21,22^).

In summary, our integrative approach assigned seven minor capsid proteins as well as the penton protein using cross-link verified AlphaFold models. This demonstrated the power of combining XL-MS, structural prediction and proteomics to elucidate the structural details of the minor and major capsid proteins, their contact sites and membrane associations.

## Discussion

Since the discovery of the first giant virus two decades ago, many new NCLDV members have been identified, expanding our understanding of the boundaries of viral complexity ^46–51^. With every newly discovered giant virus, more questions need to be addressed, urging the development of fast and systematic approaches to elucidate structure, proteomic arsenal, interaction with the hosts and ecological roles. To address this challenge, we developed an integrative approach combining XL-MS and AlphaFold3 modeling to comprehensively resolve the structural and spatial interactome of melbournevirus particles, providing unprecedented insights into its proteome organization.

Our XL-MS data partitioned the melbournevirus proteome into two spatially distinct communities separated by the inner membrane: an inner “core” and an outer “coat (Figure 1). We identified 25 TM proteins and annotated their topologies (Figure 2). Through structural assignment on the cryo-EM density map, we assigned two TM proteins (MEL_331, MEL_276) as minor capsid proteins (Figure 2 and Figure 5). These proteins likely bridge the capsid and membrane through physical contacts, which may likely explain previous observations of membrane protrusions at the capsid vertices ^22^. Notably, similar membrane-capsid tethering architecture was observed in distantly related viruses like PBCV-1 ^23^ and medusavirus ^52^, suggesting a conserved structural strategy.

Using AlphaFold3, we generated high-confidence structural models (prioritized by pTM or confidence scores and cross-link validation) and accessed functional homology with Foldseek. The coat community was enriched in proteins with predicted enzymatic functions, possibly acquired through horizontal gene transfer from the host ^53^ or other viruses ^54^. While some viral enzymes have been functionally validated ^55–57^, the activities of melbournevirus-encoded proteins remain speculative and warrant *in vitro* characterization.

Beyond enzymes, we identified a heterodimeric complex between MEL_149 and MEL_368 that forms an alternative nucleosome-like structure comparing to the previously described MEL_368:MEL_369 complex ^14^. Importantly, when MEL149:MEL_368 was modeled as a heterotetramer, nearly all cross-links from the structured regions of the MEL_149:MEL_368 complex were satisfied, further validating the model. This prediction is consistent with a recently reported *in vitro* structure of MEL_149:MEL_368 ^42^. Our data further support the existence of this complex in intact virions. However, the MEL_149:MEL_368 complex likely exists in low stoichiometry, as MEL_149 is present at fewer than 100 copies per virion, which is much lower in comparison to the ∼4,000 copies of MEL_368. Therefore, the MEL_149-containing complex may not significantly contribute to overall viral genome condensation, but it could play a regulatory role, potentially modulating transcription activation as observed in yeast ^58^.

Finally, we demonstrated the power of our integrative structural modeling approach by assigning eight additional capsid components to a previously published cryo-EM map ^45^. XL-MS data enabled us to generate a shortlist of potential protein candidates, significantly improving the success rate of matching models with the density. For regions with low-confident structural predictions or cryo-EM densities of limited resolution, the additional constraints provided by cross-linking data substantially increases assignment confidence.

We further improved our cryo-EM assignments using two complementary approaches: (i) protein copy number quantification through proteomics, and (ii) homomultimeric state predictions using AlphaFold3 ^59^. These methods successfully confirmed known assemblies, including the trimeric MCP, homodimeric MEL_213b and homopentameric MEL_256. Additionally, we predicted homo-oligomerization for putative enzymes such as MEL_219 and MEL_168, both of which share structural similarity with known enzymes of defined homo-oligomeric states ^60,61^.

While we were unable to match all remaining capsid densities to our predicted models—potentially due to protein flexibility or limitations in AlphaFold predictions— the PPI network and copy number analysis strongly implicate MEL_052, MEL_066, MEL_215, MEL_223b, MEL_232, and MEL_274 as capsid-associated proteins. These candidates represent prime targets for future structural characterization of the melbournevirus capsid.

In summary, we presented an integrative workflow combining XL-MS, proteomics, cryo-EM, and computational modeling to gain system-level insights into the spatial and structural interactome of the melbournevirus particles. Importantly, our approach only requires native virion particles, without the need of genetic manipulation. Therefore it can be easily transferred to structurally characterize other giant viruses (e.g., Marseilleviridae and NCLDV members) and other emerging pathogens, offering a powerful platform for high-throughput characterization of understudied biological systems.

## Methods

### Propagation of melbournevirus in *Acanthamoeba castellanii*

Purified melbournevirus was produced as previously described ^16,20^. In brief, *Acanthamoeba castelanii* cells were cultured in PPYG medium (2.0% w/v proteose peptone, 0.1% w/v yeast extract, 4□mM MgSO4, 0.4□mM CaCl2, 0.05□mM Fe(NH4)2(SO4)2, 2.5□mM Na2HPO4, 2.5□mM KH2PO4, and 100□mM sucrose, pH 6.5) at 28 °C in a 5% CO2 incubator. Once the cells reached 100% confluency in ten 75 cm2 cell culture flasks, each flask was infected with 10 µL of seed melbournevirus. After three days of infection, the infected culture fluid (ICF) was harvested for purifying melbournevirus particles.

### Purification of melbournevirus particles for bottom-up and cross-linking proteomics

A total of 100 mL of infected cell culture fluid was centrifuged at 600 xg, 4 °C for 15 min. The supernatant was further centrifuged at 8,500 xg, 4 °C for 35 min. The virus pellet was resuspended in 475□µL PBS, cross-linked with 5□mM freshly prepared DSSO in anhydrous DMSO, and incubated for 30□min at room temperature with shaking. Cross-linking was quenched with 15□µL of 1□M Tris (pH 8) for 30□min. The sample was rinsed once with 50 mL of 0.22-µm membrane filtered PBS and centrifuged at 8,500 xg, 4 °C for 35 min. The pellet was then resuspended in 1 mL of PBS and applied to a 10-60% w/v sucrose gradient. Centrifugation was performed at 8,500 xg, 4°C for 90 min using an ultracentrifuge (Beckman Courter, Optima XPN-100, Sw40Ti rotor). The virus band was collected and rinsed three times with 50 mL of PBS by pelleting between rinses. Finally, the pellet was thoroughly resuspended in 100 µL of lysis buffer containing 8 M Urea, 1% v/v Triton X-100, 5 mM TCEP, 50 mM TEAB (pH 8). The sample was stored at -80 °C until further processing.

### Sample preparation for bottom-up proteomics

Lysis of melbournevirus particles was facilitated by sonication in a Bioruptor pico system at 10 °C using 30s pulses of sonication, followed by 30s of pause in 10 repetitions. Genomic DNA was digested by addition of 1 µl Benzonase per sample, followed by incubation for 30 min at room temperature. Afterwards, proteins were extracted and precipitated by methanol/chloroform precipitation. Protein pellets were resuspended in a buffer containing 1% (w/v) SDC and 5 mM TCEP as well as 40 mM Iodoacetamide in TEAB. Trypsin and Lys-C were added in a protein to enzyme ratio of 1:25 and 1:100 respectively and digestion was carried out overnight at 37 °C while shaking at 1000 rpm. Next, digestion was stopped by acidification of the sample with 1% (v/v) formic acid. After centrifugation, peptides were desalted using C18 stage-tips and samples were dried by speed vacuum and stored at -20 °C until LC-MS measurement.

### Sample preparation for XL-MS

Sample preparation of cross-linked viral particles was carried out analogously to the bottom-up proteomics samples, with the exception that samples were sonicated with a Sonopuls HD 2070 Sonicator equipped with a MS73 Sonotrode (Bandelin) on 50% intensity for 30 s switching between sonication and pause every 0.5 s. After digestion, samples were desalted using c8 Sep-pak cartridges and peptide amount was determined by Pierce quantitative colorimetric assay (Thermo Fisher Scientific). After drying in speed vacuum, approximately 300 to 500 μg of peptides were loaded onto a PolySULFOETHYL A column (PolyLC) connected to an Agilent 1260 Infinity II system and peptides were separated using a 95 min gradient from buffer A (0.1% TFA in 20% ACN) to buffer B (0.5M NaCL and 0.1% TFA and 20% ACN) while collecting fractions in 60 s intervals. All fractions were then desalted individually by stage-tipping, dried in speed vacuum and stored at -20 °C before LC-MS measurement.

### LC-MS measurement

Bottom-up proteomics samples were measured on an Orbitrap Fusion Tribrid instrument operating with Tune 4.0 and Xcalibur 4.6 and online-connected to an Ultimate 3000 RSLC nano LC system (Thermo Fisher Scientific). Peptides were run over an in-house packed c18 column (column material: Poroshell 120 EC-C18, 2.7□µm; Agilent Technologies) with a flow rate of 250 nl/min using a 180 minutes gradient going from buffer A (0.1% FA in water) to buffer B (0.1% FA in 80% Acetonitrile). Instrument parameters were set as follows: MS1 Orbitrap resolution 120,000, Scan-Range 375-1500 m/z, AGC-target “standard”, maximum injection time 50 ms. Duty cycles were set to 1 s. Intensity threshold for precursors was set to 1.0E+4. Precursors of Charge 2-4 were selected with dynamic exclusion of 40 s and fragmented by Higher collisional energy dissociation (HCD) with normalized collision energy set to 30%. MS2 scans were recorded in the Ion Trap with scan rate set to “rapid” and scan range set to “auto”. AGC-target for MS2 was set to “standard”. Cross-linking proteomics samples were measured on an Orbitrap Fusion Lumos mass spectrometer (Thermo Fisher Scientific) equipped with a FAIMS Pro Duo interface (Thermo Fisher Scientific) operating with Tune 4.0 and Xcalibur 4.6. Separation was performed on an in-house packed C18 column using the same LC gradient and flow as for bottom-up samples. FAIMS compensation voltages were set to -50, -60 and -75 V. Instrument parameters were set as follows: MS1 Orbitrap resolution 120,000, Scan-Range 375-1600 m/z, AGC-target “standard”, maximum injection time 50 ms. Duty cycles were set to 2 s. Intensity threshold for precursors was set to 2.0E+4. Charge state filter for precursor selection was set to 4-8 and dynamic exclusion set to 60 s. Precursors were fragmented by stepped-HCD with normalized collision energy set to 21%, 27% and 33%. MS2 scans were recorded in the Orbitrap with resolution set to 60,000 and scan range set to “auto”. AGC-target for MS2 was set to 200% and maximum injection time was set to 118 ms.

### Bottom-up proteomics data analysis

Raw files were searched with MaxQuant version 2.6.7.0 ^62^ with standard parameters, match between runs and iBAQ enabled. Methionine oxidation and N-terminus acetylation were set as variable modifications, carbamidomethylation of cysteine was set as fixed modification. Trypsin was selected as a digestion enzyme. Precursor mass tolerance was set to 20 ppm and fragment mass tolerance was set to 0.5 Da. Spectra were searched against the melbournevirus proteome database (TaxonID: 1560514 ^16^) as well as a database of the host Acanthamoeba castellanii previously described to be assembled from the reference database plus sequences received by RNA sequencing ^33^. Results were filtered to 1% FDR at PSM and protein level.

For targeted search for protein ubiquitination, the bottom-up proteomics rawfiles were analysed with Fragpipe (v22.0) MSFragger using the standard ubiquitin modification workflow (K, +114.04293 Da)^63,64^.

### Copy number estimation of virion incorporated proteins

Calculation of copy numbers was conducted in R as described before ^29^. Protein iBAQ-values from the proteinGroups.txt file were used for quantification. Proteins MEL_305, MEL_236, MEL_213b and MEL_256 were used as internal standards with known copy numbers obtained from previous findings of cryo-EM (MEL_305: 9240 copies, MEL_236: 9240 copies, MEL_213b: 120 copies, MEL_256: 60 copies). For each of the 4 technical replicates a linear regression of log10 iBAQ values against log10 absolute copy numbers per virion was calculated. Copy numbers of remaining proteins were calculated using the slope and offset of the individual regression grade following the equation:

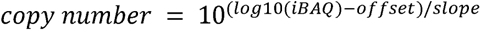

The mean of the calculated copy numbers from the 4 replicates was used.

### XL-MS data analysis

Rawfiles of XL-MS samples were searched using Scout software (version 1.5.1) ^65^ against a database assembled from all melbournevirus proteins and all host proteins identified in the bottom-up proteomics experiments. The following parameters were selected: MS1 tolerance 10 ppm, MS2 tolerance 20 ppm, minimum peptide length 6 amino acids, maximum peptide length 60 amino acids, 3 missed cleavages and 2 variable modifications. Cross-linker type pre-settings for DSSO were used (158.00376533 Da, short arm 54.01056468 Da, long arm 85.98263585 Da) with residues K, S, T, Y and protein N-terminus as possible cross-linking sites. Results were reported at a 1% FDR at residue-pair-level.

### Transmembrane domain prediction

All transmembrane domain prediction software were installed and run locally on Ubuntu 22.04.1 LTS using the melbournevirus protein sequences from the Uniprot database as input files. DeepTMHMM was run using pybiolib package with standard workflow described at https://dtu.biolib.com/DeepTMHMM. Probability files were used as results. DeepTMpred was run with python using model files supplied at https://github.com/ISYSLAB-HUST/DeepTMpred. Probabilities were extracted from JSON result files. TMbed was run using python with standard parameters as described at https://github.com/BernhoferM/TMbed. Results were generated in output-format 3 and probabilities were extracted from .pred files.

### Structure prediction with AlphaFold

AlphaFold version 3.0.1 ^5^ was installed and run locally on Ubuntu 24.04.1 LTS by utilizing the Docker platform. Predictions were run on a NVIDIA RTX A6000 GPU (48 GB RAM). Number of recycles was set to default. We used the initial AlphaFold3 input format, the default parameters as well as databases supplied at https://github.com/google-deepmind/alphafold3. Model parameters were received directly from Google. Number of multimer predictions per model was set to 1. The heteromultimeric model of MEL_149 and MEL_368 has been predicted with the AlphaFold3 online search server. pTM and ipTM scores were extracted from .json files using R and PAE data was extracted using python 3. For every monomer or multimer prediction the model with the best agreement with cross-linking data was selected. In case of equally satisfied cross-links, the model with highest pTM, or in case of multimeric predictions ipTM score were selected for analysis.

Search for structurally related proteins was conducted using Foldseek ^44^. This algorithm returns characterized protein structures from PDB and also the AlphaFold database that share structural similarity to the input model. The quality of the alignment was addressed on the basis of the given scores (e.g. TM-score or probability score).

### Interpretation of cryo-EM data

For comparison of predicted models with cryo-EM data the 3.4 □ resolution map of the melbournevirus capsid vertex described in ^45^ has been loaded in UCSF ChimeraX v1.8 ^66^. Pictures were taken at volume step 1 and threshold level 0.012. Predicted AlphaFold3 models were loaded and fitted into the map using the “fit in map” tool. For predicted models with lower-confidence parts only the well-predicted parts were used for the fitting.

For refinement of AlphaFold models fitted into the global capsid density map, the corresponding local area of the map was masked and saved as a new volume. For each of these new maps, the target density within the mask region was shifted to the center of the box followed by resampling the map to a smaller box size. Each model was then rigid-body docked into the resampled map and refined in ISOLDE ^67^ with distance and angular restraints enabled. Some stretches of the models were manually tacked into the density and rebuilt. Residues for which no matching density could be found were deleted. Adjusted models were then real space refined against the corresponding maps in phenix ^68^ with default parameters and secondary structure restraints enabled. For the MCP trimer and pentamer models’ symmetry constraints and refinement of symmetry operators were included as well.

## Data availability

All rawfiles corresponding to this manuscript have been uploaded to the ProteomeXchange Consortium via the PRIDE ^69^ partner repository with the dataset identifier PXD064933. AlphaFold models have been deposited at Figshare under 10.6084/m9.figshare.29290655.

## Acknowledgements

The authors thank Prof. Chantal Abergel for supplying the seed virus for melbournevirus cultivation. FL and LM are supported by DFG Project LI 3260/6-1 and Leibniz-Wettbewerb (P70/2018). JR is funded by European Research Council (ERC) Starting Grant (ERC-STG No. 949184). KO is funded by the Swedish Research Council (grant number: 2018-03387 and 2023-01857) and the Carl Tryggers Stiftelse (grant number: CTS23:2703). KM is funded by AMED BINDS (grant number: 24ama121005j0003). MK is supported by the Helmholtz Society and the Heisenberg Professorship from DFG (KU 3222/3-1). The authors thank Prof. Chantal Abergel for critical reading of the manuscript, Dr. Libera Lo Presti and Dr. Philip Lössl for editing the manuscript. The cryo-EM data was collected at the Cryo-EM Swedish National Facility funded by the Knut and Alice Wallenberg, Erling Persson Family, and Kempe Foundations, SciLifeLab, Stockholm University and Umeå University, and processed at ExCELLS/NIPS, NINS, Japan.

## Author contributions

LM, BB, KO and FL conceptualized the project. LM and KO performed experiments. LM analyzed experimental data. JR performed AlphaFold prediction. VM and MK refined protein models according to cryo-EM data. RBS and KM supplied access to cryo-EM data. LM created figures and wrote the original draft. BB and FL reviewed and edited the manuscript. All authors revised the manuscript.

## Competing Interest

FL is a shareholder and advisory board member of Absea Biotechnology and VantAI. The remaining authors declare no competing interests.

---

**Supplementary Figure 1.**
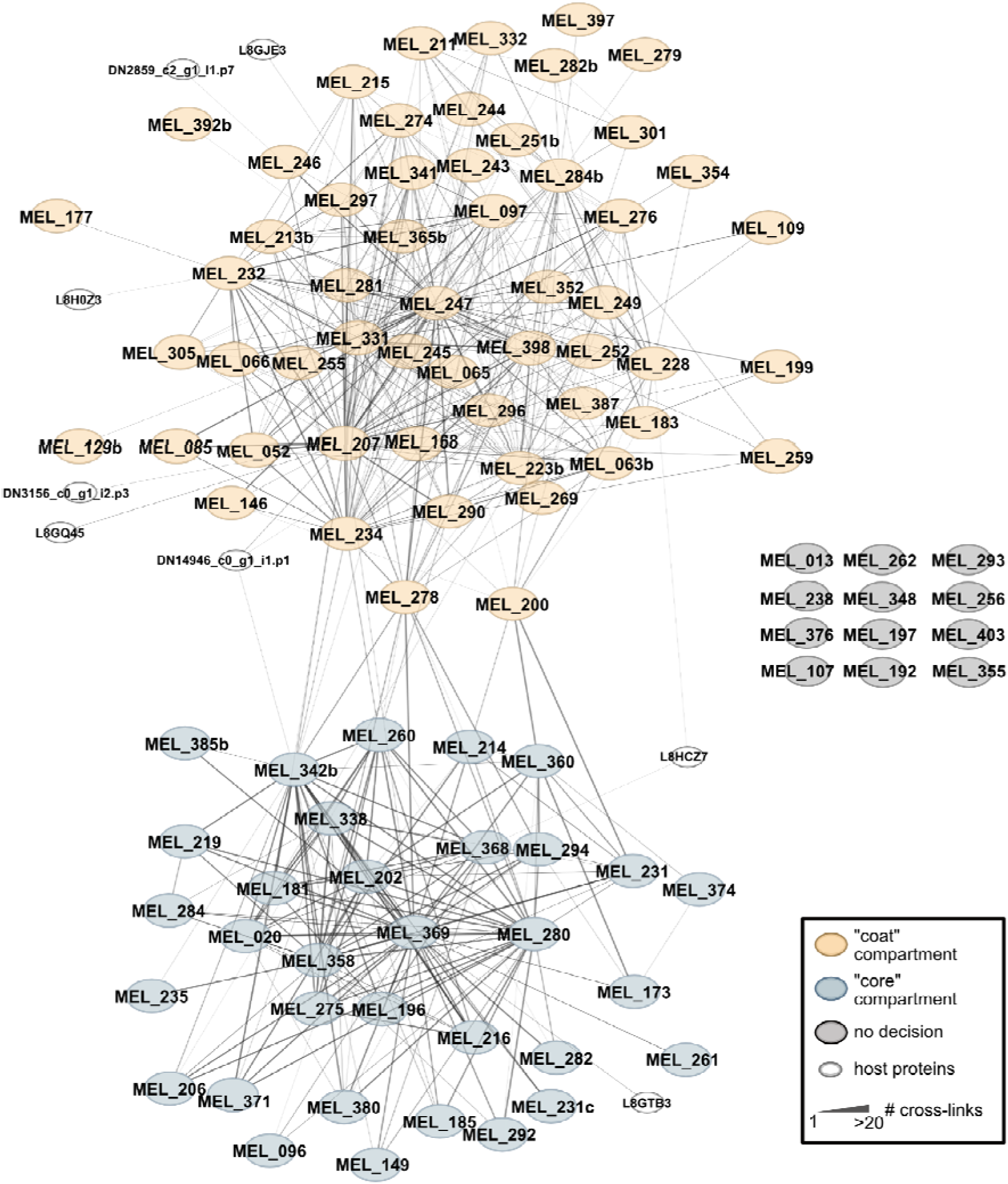
Extended view of the melbournevirus protein interactome. XL-MS-based interaction network of melbournevirus as presented in Figure 1C. Protein gene names are annotated corresponding to the gene names from the Uniprot database. For host proteins lacking assigned gene names, Uniprot identifiers or transcriptomics-derived database annotations were used instead.

**Supplementary Figure 2.**
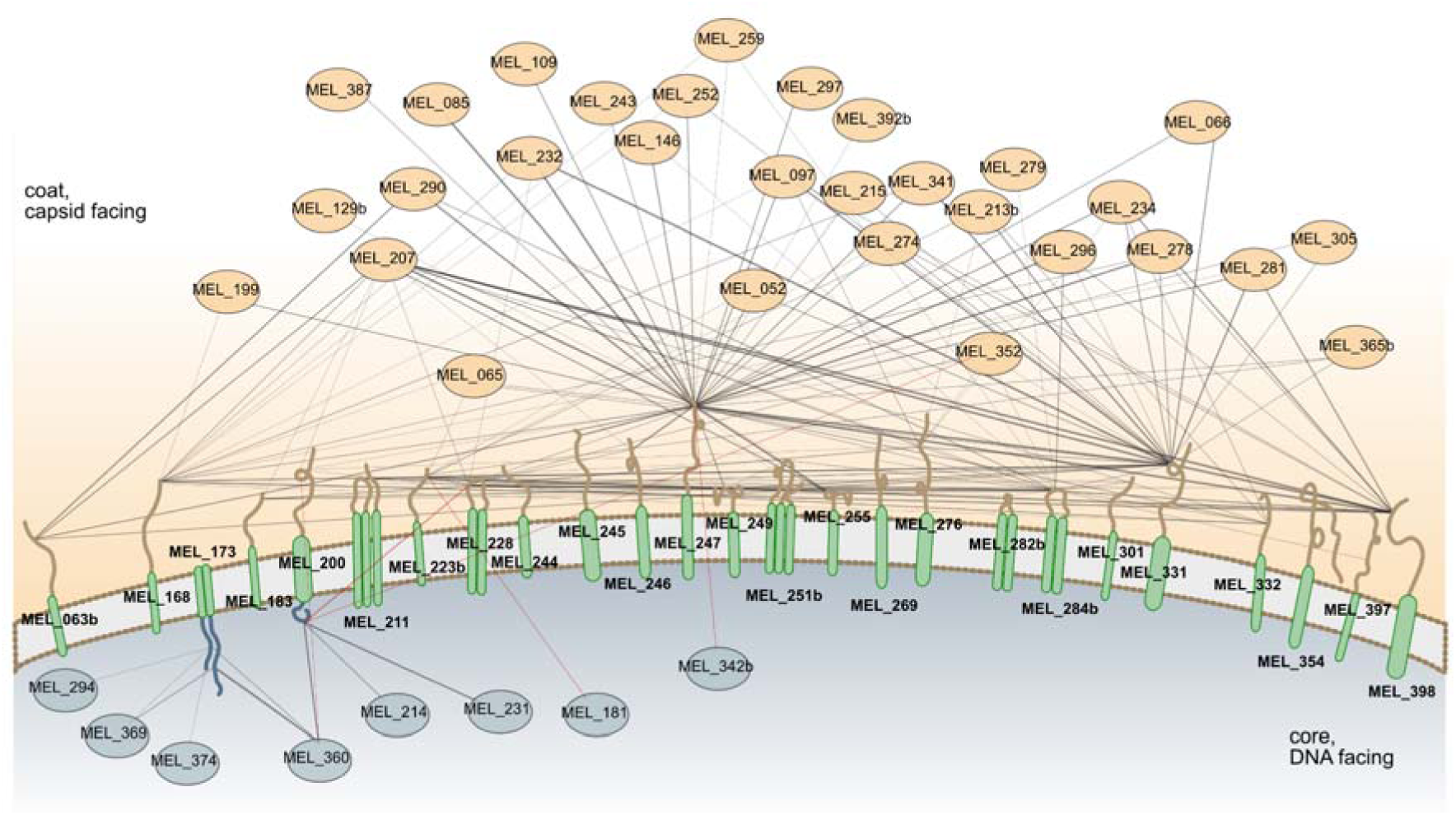
Extended view of the membrane-proximal protein interactome of predicted membrane proteins. Overview of predicted transmembrane proteins, their topologies and their direct interaction partners, with all gene names annotated. Proteins are color-coded as in Figure 2B and 2C. For clarity, cross-links within the same domain targeting the same interactor are grouped.

**Supplementary Figure 3.**
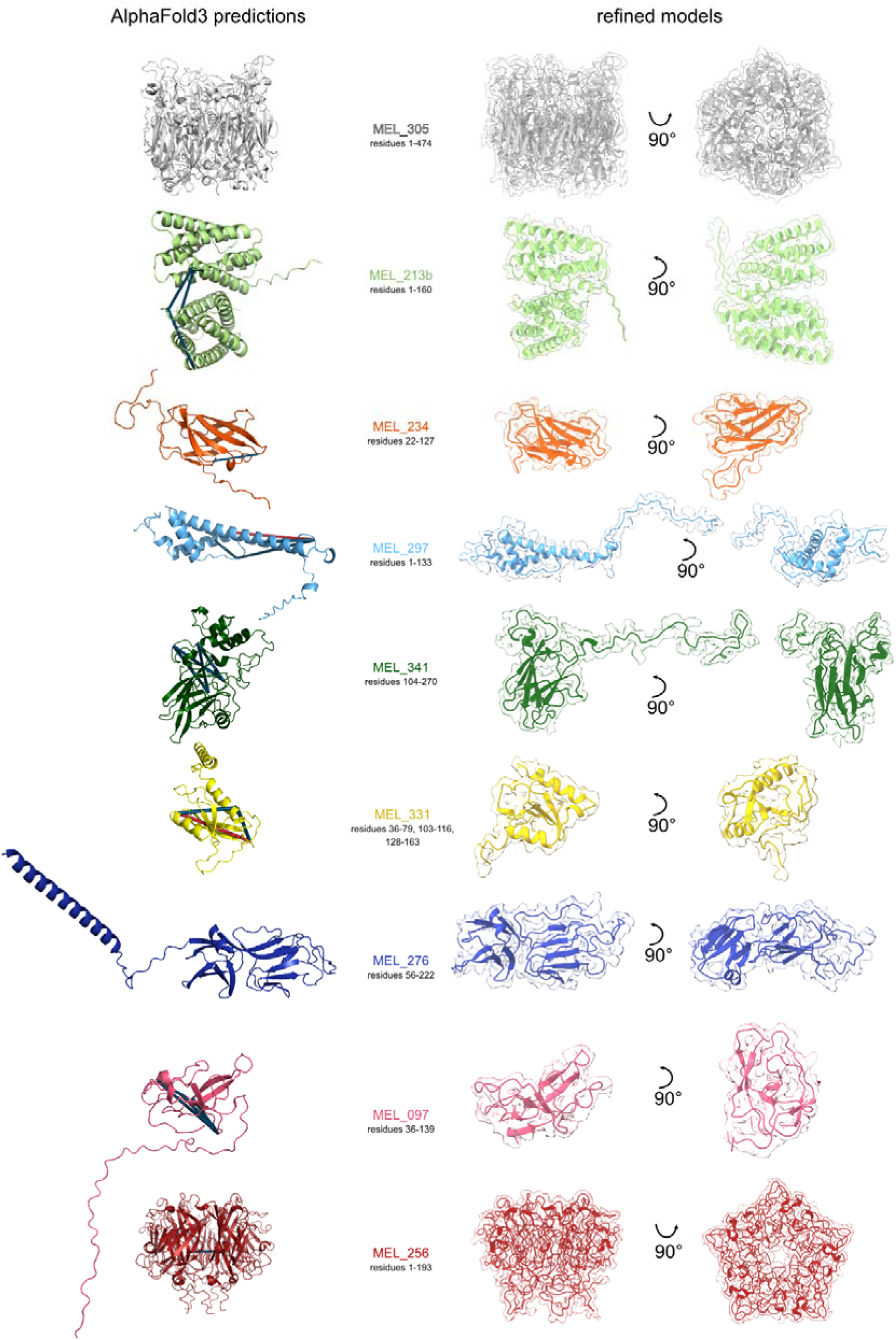
Fitting refined AlphaFold models into cryo-EM density. AlphaFold3 models the MCP (MEL_305), minor capsid proteins and penton protein in their assigned stoichiometry in this study, with mapped intra-protein cross-links (left). Cross-links satisfying the DSSO distance constraint (≤35 Å, Cα–Cα) are shown in blue; over-length cross-links are in red. MEL_305 has no intra-link detected. MEL_276 only has intra-links from disordered regions. Cryo-EM density-guided refinement of models (right). Regions that could be traced are indicated below each protein name.

**Supplementary Figure 4.**
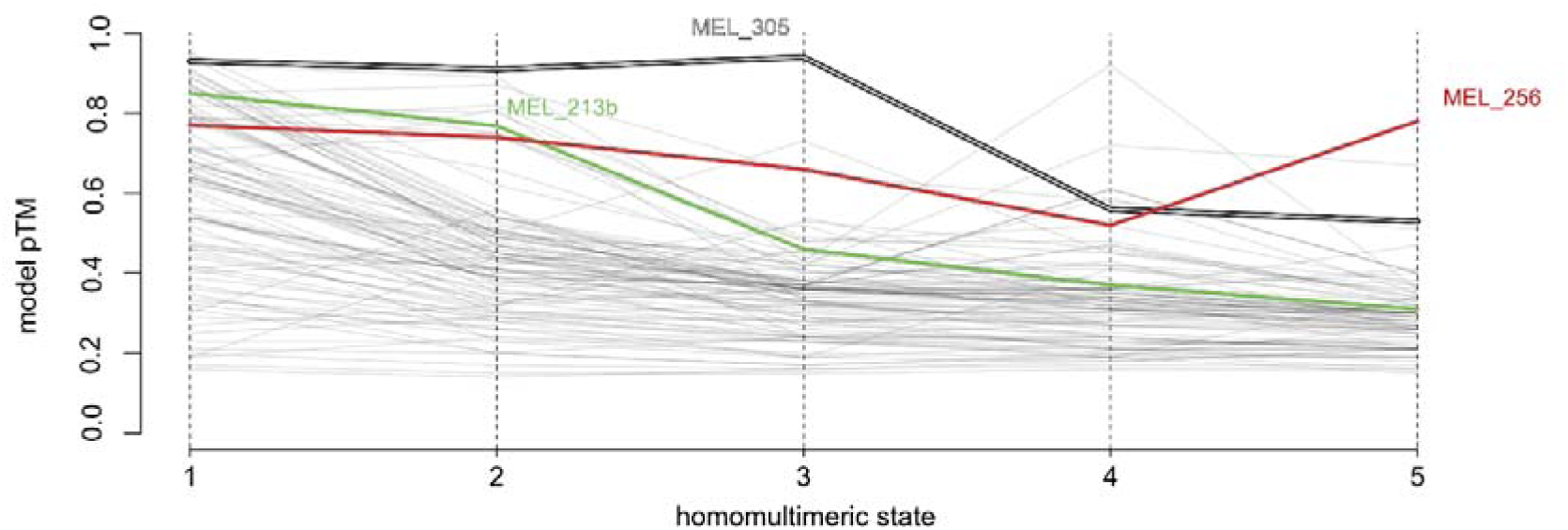
Prediction of protein multimeric states using AlphaFold confidence scores. Homomultimeric states were predicted for all proteins identified in the XL-MS dataset. The local maximum of the pTM score was used to determine the most likely oligomeric state. Two proteins - MEL_305 (MCP) and MEL_213b - with previously known stoichiometry as well as MEL_256 - the hereby identified penton protein - are highlighted.

**Supplementary Figure 5.**
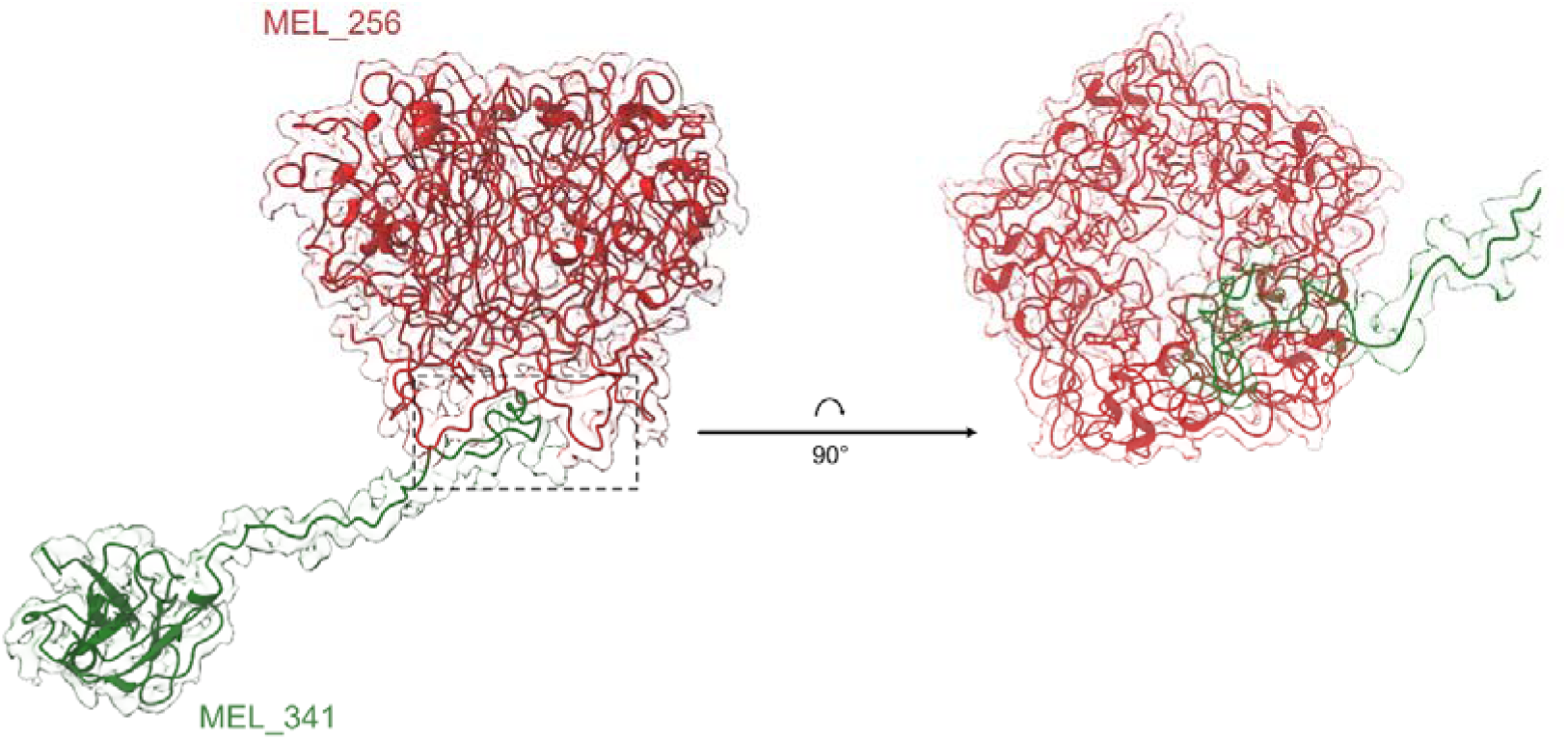
Cryo-EM-guided structural refinement reveals interaction between MEL_341 and the penton protein. Refined AlphaFold models of pentameric MEL_256 (red) and MEL_341 (green) fitted into the cryo-EM density. A flexible region of MEL_341 (highlighted) mediates direct contact with the penton protein beneath the capsid vertex.

**Supplementary Figure 6.**
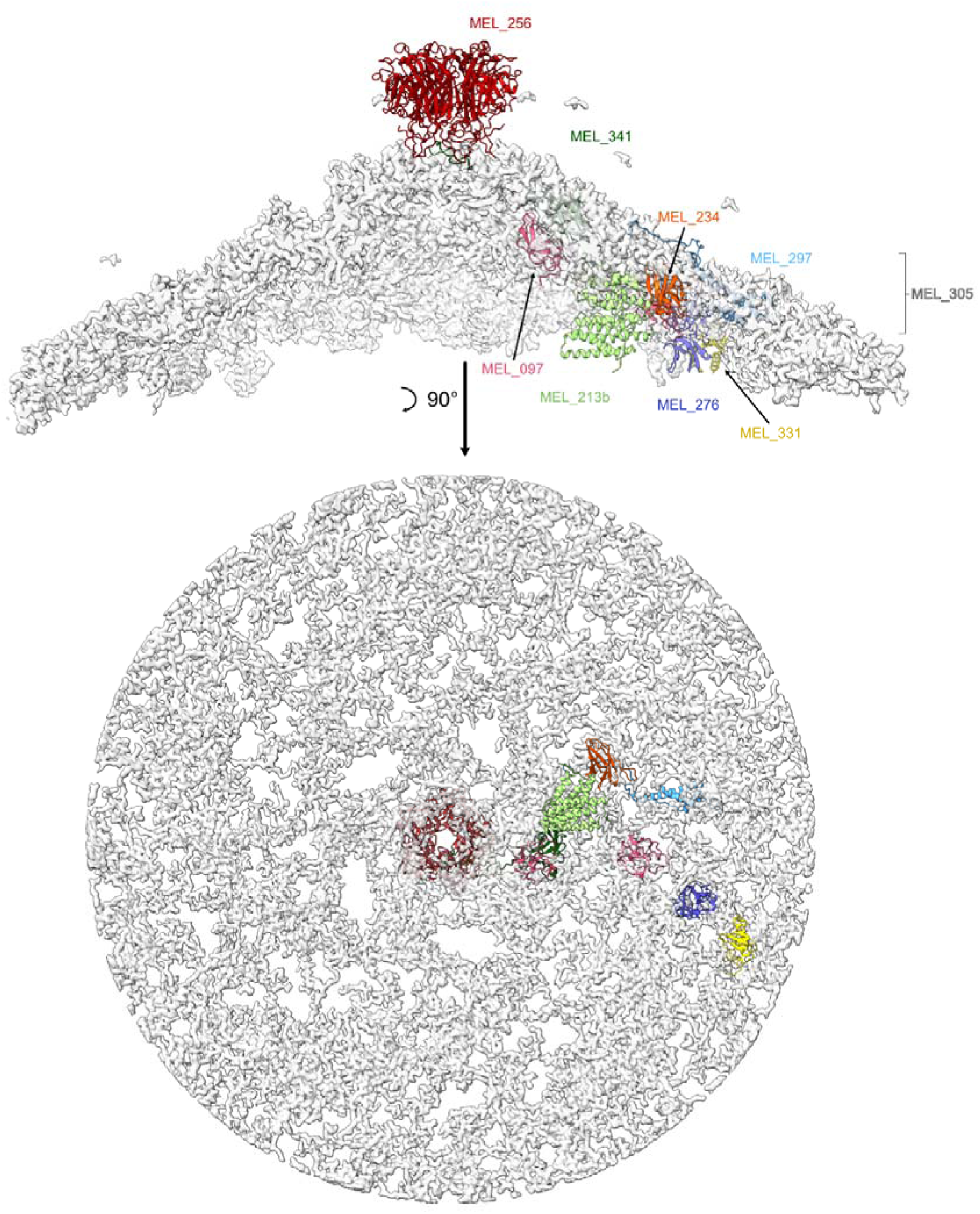
Unassigned density in melbournevirus capsid vertex. Five-fold symmetric capsid vertex shown in side and interior views. Previously identified and newly assigned proteins are highlighted in different colors. Density around these identified components is removed. Remaining density is shown in gray.

**Supplementary Figure 7.**
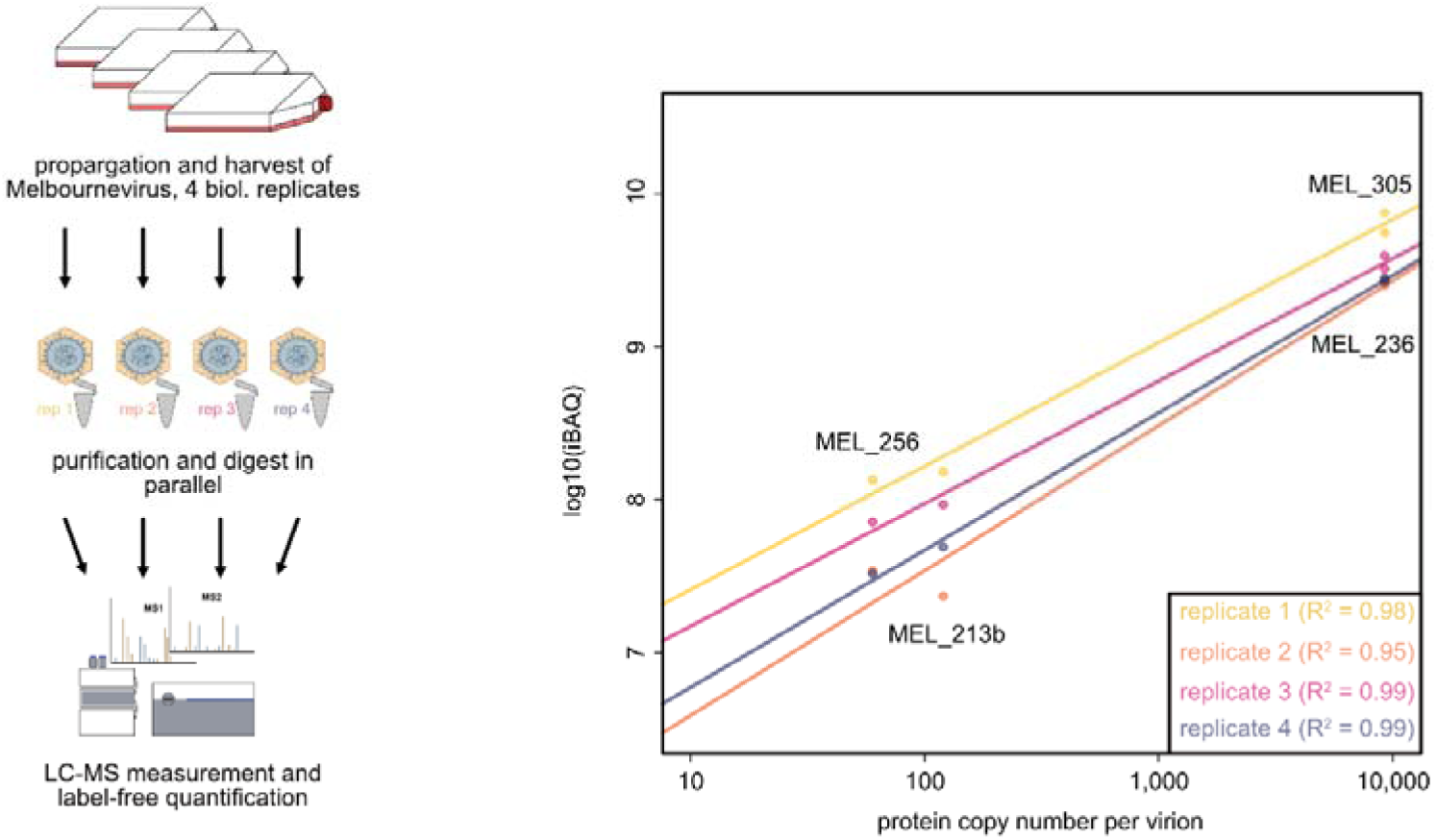
Quantitative proteomics-based estimation of protein copy numbers. Protein iBAQ values were determined by label-free bottom-up proteomics across four technical replicates. Theoretical copy numbers were derived by plotting known capsid protein stoichiometries on a double-logarithmic scale. The linear relationship (reference lines shown) enables extrapolation of copy numbers for all other viral proteins based on their iBAQ values.

## Reference list

1. Hambly, E. & Suttle, C.A. The viriosphere, diversity, and genetic exchange within phage communities. Current Opinion in Microbiology 8, 444–450 (2005).

2. Suttle, C.A. Viruses in the sea. Nature 437, 356–361 (2005).

3. Suttle, C.A. Viruses: a vast reservoir of genetic diversity and driver of global processes. Retrovirology 6, I7 (2009).

4. Jumper, J. et al. Highly accurate protein structure prediction with AlphaFold. Nature 596, 583–589 (2021).

5. Abramson, J. et al. Accurate structure prediction of biomolecular interactions with AlphaFold 3. Nature 630, 493–500 (2024).

6. Nomburg, J. et al. Birth of protein folds and functions in the virome. Nature 633, 710–717 (2024).

7. Gomes, P.S.F.C., Gomes, D.E.B. & Bernardi, R.C. Protein structure prediction in the era of AI: Challenges and limitations when applying to in silico force spectroscopy. Frontiers in Bioinformatics 2(2022).

8. Perdigão, N. et al. Unexpected features of the dark proteome. Proceedings of the National Academy of Sciences 112, 15898–15903 (2015).

9. Colson, P. et al. “Megavirales”, a proposed new order for eukaryotic nucleocytoplasmic large DNA viruses. Archives of Virology 158, 2517–2521 (2013).

10. Abergel, C., Legendre, M. & Claverie, J.M. The rapidly expanding universe of giant viruses: Mimivirus, Pandoravirus, Pithovirus and Mollivirus. Fems Microbiology Reviews 39, 779–796 (2015).

11. Philippe, N. et al. Pandoraviruses: Amoeba Viruses with Genomes Up to 2.5 Mb Reaching That of Parasitic Eukaryotes. Science 341, 281–286 (2013).

12. La Scola, B. et al. A giant virus in amoebae. Science 299, 2033–2033 (2003).

13. Raoult, D. et al. The 1.2-megabase genome sequence of mimivirus. Science 306, 1344–1350 (2004).

14. Liu, Y. et al. Virus-encoded histone doublets are essential and form nucleosome-like structures. Cell 184, 4237–4250.e19 (2021).

15. Talbert, P.B., Armache, K.-J. & Henikoff, S. Viral histones: pickpocket’s prize or primordial progenitor? Epigenetics & Chromatin 15(2022).

16. Doutre, G., Philippe, N., Abergel, C. & Claverie, J.-M. Genome Analysis of the First Marseilleviridae Representative from Australia Indicates that Most of Its Genes Contribute to Virus Fitness. Journal of Virology 88, 14340–14349 (2014).

17. Abergel, C. & Claverie, J.-M. Giant viruses. Current Biology 30, R1108–R1110 (2020).

18. Thomas, V. et al. Lausannevirus, a giant amoebal virus encoding histone doublets. Environmental Microbiology 13, 1454–1466 (2011).

19. Arantes, T.S. et al. The Large Marseillevirus Explores Different Entry Pathways by Forming Giant Infectious Vesicles. Journal of Virology 90, 5246–5255 (2016).

20. Okamoto, K. et al. Cryo-EM structure of a Marseilleviridae virus particle reveals a large internal microassembly. Virology 516, 239–245 (2018).

21. Burton-Smith, R.N. et al. The 4.4 Å structure of the giant Melbournevirus virion belonging to the Marseilleviridae family. (bioRxiv - Cold Spring Harbor Laboratory, 2021).

22. Chihara, A. et al. A novel capsid protein network allows the characteristic internal membrane structure of Marseilleviridae giant viruses. Scientific Reports 12(2022).

23. Fang, Q. et al. Near-atomic structure of a giant virus. Nature Communications 10(2019).

24. Shao, Q. et al. Near-atomic, non-icosahedrally averaged structure of giant virus Paramecium bursaria chlorella virus 1. Nature Communications 13(2022).

25. Zhang, X. et al. Three-dimensional structure and function of the Paramecium bursaria chlorella virus capsid. Proceedings of the National Academy of Sciences 108, 14837–14842 (2011).

26. Bryson, T.D. et al. A giant virus genome is densely packaged by stable nucleosomes within virions. Molecular Cell 82, 4458-+ (2022).

27. Liu, F., Rijkers, D.T.S., Post, H. & Heck, A.J.R. Proteome-wide profiling of protein assemblies by cross-linking mass spectrometry. Nature Methods 12, 1179–1184 (2015).

28. Zhu, Y. et al. Cross-link assisted spatial proteomics to map sub-organelle proteomes and membrane protein topologies. Nature Communications 15(2024).

29. Bogdanow, B. et al. Spatially resolved protein map of intact human cytomegalovirus virions. Nature Microbiology 8, 1732–1747 (2023).

30. Bogdanow, B. et al. Redesigning error control in cross-linking mass spectrometry enables more robust and sensitive protein-protein interaction studies. Molecular Systems Biology 21, 90–106 (2025).

31. Bartolec, T.K. et al. Cross-linking mass spectrometry discovers, evaluates, and corroborates structures and protein–protein interactions in the human cell. Proceedings of the National Academy of Sciences 120(2023).

32. O’Reilly, F.J. et al. Protein complexes in cells by AI-assisted structural proteomics. Molecular Systems Biology 19(2023).

33. Bernard, C. et al. A time-resolved multi-omics atlas of Acanthamoeba castellanii encystment. Nature Communications 13(2022).

34. Fabre, E. et al. Noumeavirus replication relies on a transient remote control of the host nucleus. Nature Communications 8, 15087 (2017).

35. Girvan, M. & Newman, M.E.J. Community structure in social and biological networks. Proceedings of the National Academy of Sciences 99, 7821–7826 (2002).

36. Hallgren, J. et al. DeepTMHMM predicts alpha and beta transmembrane proteins using deep neural networks. (bioRxiv - Cold Spring Harbor Laboratory, 2022).

37. Wang, L., Zhong, H., Xue, Z. & Wang, Y. Improving the topology prediction of alpha-helical transmembrane proteins with deep transfer learning. Comput Struct Biotechnol J 20, 1993–2000 (2022).

38. Bernhofer, M. & Rost, B. TMbed: transmembrane proteins predicted through language model embeddings. BMC Bioinformatics 23(2022).

39. Sun, J.F. et al. Machine learning in computational modelling of membrane protein sequences and structures: From methodologies to applications. Computational and Structural Biotechnology Journal 21, 1205–1226 (2023).

40. Chen, Y.-J. et al. X-ray structure of EmrE supports dual topology model. Proceedings of the National Academy of Sciences 104, 18999–19004 (2007).

41. Wang, X.R. et al. Molecular Details Underlying Dynamic Structures and Regulation of the Human 26S Proteasome. Molecular & Cellular Proteomics 16, 840–854 (2017).

42. Villalta, A., Bisio, H., Toner, C.M., Abergel, C. & Luger, K. Melbournevirus encodes a shorter H2B-H2A doublet histone variant that forms structurally distinct nucleosome structures. (Cold Spring Harbor Laboratory, 2025).

43. Varadi, M. et al. AlphaFold Protein Structure Database in 2024: providing structure coverage for over 214 million protein sequences. Nucleic Acids Research (2023).

44. Van Kempen, M. et al. Fast and accurate protein structure search with Foldseek. Nature Biotechnology 42, 243–246 (2024).

45. Burton-Smith, R.N. & Murata, K. Post-acquisition super resolution for cryo-electron microscopy. (bioRxiv - Cold Spring Harbor Laboratory, 2023).

46. Schulz, F. et al. Giant viruses with an expanded complement of translation system components. Science 356, 82–85 (2017).

47. Yoshikawa, G. et al. Medusavirus, a Novel Large DNA Virus Discovered from Hot Spring Water. Journal of Virology 93(2019).

48. Andreani, J. et al. Pacmanvirus, a New Giant Icosahedral Virus at the Crossroads between Asfarviridae and Faustoviruses. Journal of Virology 91, JVI.00212-17 (2017).

49. Arthofer, P. et al. A giant virus infecting the amoeboflagellate Naegleria. Nature Communications 15(2024).

50. Schulz, F. et al. Hidden diversity of soil giant viruses. Nature Communications 9(2018).

51. Sheng, Y.J., Wu, Z.Q., Xu, S.Z. & Wang, Y.J. Isolation and Identification of a Large Green Alga Virus (Virus XW01) of Mimiviridae and Its Virophage (Virus Virophage SW01) by Using Unicellular Green Algal Cultures. Journal of Virology 96(2022).

52. Watanabe, R., Song, C., Takemura, M. & Murata, K. Subnanometer structure of medusavirus capsid during maturation using cryo-electron microscopy. Journal of Virology 98(2024).

53. Filée, J. & Chandler, M. Gene Exchange and the Origin of Giant Viruses. Intervirology 53, 354–361 (2010).

54. Wu, J.Y. et al. Gene Transfer Among Viruses Substantially Contributes to Gene Gain of Giant Viruses. Molecular Biology and Evolution 41(2024).

55. Antika, T.R. et al. A naturally occurring mini-alanyl-tRNA synthetase. Communications Biology 6(2023).

56. Benarroch, D., Claverie, J.-M., Raoult, D. & Shuman, S. Characterization of Mimivirus DNA Topoisomerase IB Suggests Horizontal Gene Transfer between Eukaryal Viruses and Bacteria. Journal of Virology 80, 314–321 (2006).

57. Piacente, F. et al. Characterization of a UDP--acetylglucosamine biosynthetic pathway encoded by the giant DNA virus Mimivirus. Glycobiology 24, 51–61 (2014).

58. Raisner, R.M. et al. Histone Variant H2A.Z Marks the 5′ Ends of Both Active and Inactive Genes in Euchromatin. Cell 123, 233–248 (2005).

59. Akdel, M. et al. A structural biology community assessment of AlphaFold2 applications. Nature Structural & Molecular Biology 29, 1056-+ (2022).

60. Ollis, D.L. et al. The Alpha/Beta-Hydrolase Fold. Protein Engineering 5, 197–211 (1992).

61. Nicholson, A.W. Ribonuclease III mechanisms of double□stranded RNA cleavage. WIREs RNA 5, 31–48 (2014).

62. Cox, J. & Mann, M. MaxQuant enables high peptide identification rates, individualized p.p.b.-range mass accuracies and proteome-wide protein quantification. Nature Biotechnology 26, 1367–1372 (2008).

63. Kong, A.T., Leprevost, F.V., Avtonomov, D.M., Mellacheruvu, D. & Nesvizhskii, A.I. MSFragger: ultrafast and comprehensive peptide identification in mass spectrometry–based proteomics. Nature Methods 14, 513–520 (2017).

64. Cifani, P. et al. Discovery of Protein Modifications Using Differential Tandem Mass Spectrometry Proteomics. Journal of Proteome Research 20, 1835–1848 (2021).

65. Clasen, M.A. et al. Proteome-scale recombinant standards and a robust high-speed search engine to advance cross-linking MS-based interactomics. Nature Methods 21, 2327–2335 (2024).

66. Pettersen, E.F. et al. UCSF ChimeraX: Structure visualization for researchers, educators, and developers. Protein Science 30, 70–82 (2021).

67. Croll, T.I. ISOLDE: a physically realistic environment for model building into low-resolution electron-density maps. Acta Crystallographica Section D Structural Biology 74, 519–530 (2018).

68. Afonine, P.V., Headd, J.J., T.C., T. & Adams, P.D. Computational Crystallography Newsletter. Volume 4, Part 2, 43–44. edn (2013).

69. Perez-Riverol, Y. et al. The PRIDE database resources in 2022: a hub for mass spectrometry-based proteomics evidences. Nucleic Acids Research 50, D543–D552 (2022).

70. Pruneda, N., Jonathan et al. The Molecular Basis for Ubiquitin and Ubiquitin-like Specificities in Bacterial Effector Proteases. Molecular Cell 63, 261–276 (2016).

